# Measuring conscious tactile perception without report: pupil, blink, and microsaccade signatures of a tactile no-report paradigm

**DOI:** 10.64898/2026.10.29.762884

**Authors:** Ayushe Sharma, Ritvik Senjalia, Annabel Woodworth, David Jin, Shanae Aerts, Devon Cormier, Aya Khalaf, Hal Blumenfeld

## Abstract

Conscious perception is often inferred from behavioral reports, which conflate awareness-related neural signals with the motor and decision processes required to respond. No-report paradigms address this by using spontaneous physiological signals, such as eye movements, to infer perception. Although our group developed a no-report visual paradigm, we had not applied this approach to touch. We developed a no-report tactile paradigm using the Galileo stimulation system to deliver pneumatic air puffs to non-thumb fingers on both hands while tracking eye movements. Participants reported sensations from the Report hand while receiving threshold-level puffs on both hands; eye metrics were used to infer perception in the No-Report hand. Perceived threshold-level stimuli elicited greater pupil dilation and more post-stimulus blinking than identical unperceived stimuli, with effects lasting 1–2 seconds. Microsaccade rate decreased after stimulus onset, but this suppression did not reach statistical significance. A stacked support vector machine trained on the Report eye-tracking measures was applied to classify No-Report trials and successfully separated [Predicted] Perceived from [Predicted] Not Perceived trials (AUC = .92; cross-validated accuracy = 72%; sensitivity = 90% at 80% specificity; label-permutation p < .001). These findings show that eye tracking can detect conscious tactile perception without overt responses and offer a noninvasive, scalable approach for studying tactile perception in patients with self-reported disabilities, including comatose or intubated patients.

## Introduction

Traditionally, conscious perception has been operationally defined by behavioral report: if a participant responds by pressing the "yes" button, the stimulus has been perceived. While this definition is appealing, it confounds the neural signatures of awareness with processes involved in reporting them: a decision to respond (and thus the motivation or intentionality), preparation and execution of an act (a motor response), and introspection on one’s perceptual state itself.^1^ Therefore, report-based brain maps are contaminated by both conscious access as well as decisions and actions, so it is impossible to separate what may be specific to awareness alone from all these other processes that we require in order to report them.^2^ The no-report paradigm solves this problem because the physical stimulus remains constant but instead of asking for a response or reporting back on what was perceived, perception itself can now be inferred by observing spontaneous physiological signals rather than relying upon an overt behavioral one.^2^ Using such visual no-report paradigms, it has recently been demonstrated that near-threshold stimuli elicit BOLD responses in widespread cortical and subcortical areas even when there is no need to report the percept (i.e., what was seen) and single-trial measures of conscious states can be decoded from features of eye tracking alone.^3^

In our laboratory, EEG and simultaneous eye tracking in a report-based tactile paradigm have shown that perceived tactile stimuli, compared to physically identical non-perceived stimuli, elicit significantly higher N80, N140 and P300 event-related potentials and reduced microsaccade rate.^4^ In general, three properties make tactile media especially suitable for the study of consciousness: (1) touch is the first sense to develop, with peripheral mechanoreceptors being functional by the seventh week of gestation, and tactile responsiveness documented in the fetus long before visual and auditory perception,^5,6^ thus making touch the earliest neural substrate for consciousness; (2) touch is a window into the dysfunction of consciousness,^7,8^ and (3) sensory nerve action potentials from mechanoreceptors in the skin are conducted in the dorsal column, synapse in the ventral posterolateral nucleus of the thalamus, and project to the primary somatosensory cortex^9^ — the very area of the brain damaged by stroke, trauma, and hypoxia, a thin, gyrus-like strip of the postcentral gyrus with finger representations at the central sulcus,^10^ and thus prone to partial-volume averaging and motion-related artifacting of the perceptual activity it measures. A tactile no-report design is thus not only elegant, but methodologically necessary.

The Detect-Pulse-Switch-Wave (DPSW) model of consciousness suggests a four-stage sequence of cortical and subcortical events: Detect (0 to 100 ms; the stimulus is registered by early sensory cortex and superior colliculus), Pulse (100 to 300 ms; a transient subcortical arousal burst from the thalamus, brainstem, and basal forebrain enhances the sensory signal), Switch (200 to 500 ms; the default mode network is inhibited and attention is reallocated via basal ganglia circuits), and Wave (300 to 1000 ms; a traveling cortical activation across the frontoparietal task-positive network enables report and memory).^11^ The DPSW model predicts that the Pulse and Switch components are generated subcortically and thus should be observable in physiological signals linked to the arousal and attention systems that underlie them, even in the absence of explicit report. These signals can be measured non-invasively from a single infrared eye tracker: task-evoked pupil dilation, which indexes locus coeruleus noradrenergic activity on a 1 to 2 s latency, and expected to track the Pulse,^12,13^ spontaneous blink timing, modulated by basal ganglia dopaminergic activity on a similar timescale, and expected to track the Switch;^14,15^ and microsaccade rate, controlled by the superior colliculus, and expected to be suppressed during focused processing of a detected stimulus, tracking the Detect phase.^16^

We developed a tactile analogue of the visual no-report paradigm to test whether these three eye-based signatures of DPSW (H1, Pulse; H2, Switch; H3, Detect) are present for near-threshold tactile stimuli, generalize from Report to No-Report trials, and whether their joint information suffices to decode conscious perception on a single-trial basis without any behavioral response. We hypothesized that perceived threshold stimuli would elicit a transient pupil dilation (H1, Pulse), an increase in blink rate (H2, Switch), and a suppression of microsaccade rate (H3, Detect) relative to physically identical not-perceived stimuli, and that these signatures would be present on both Report and No-Report trials. Here, we report the behavioral and eye-tracking results.

## Materials and Methods

### Participants

We acquired 37 datasets from nineteen participants (each visited for two sessions within 2 weeks of the first session except one participant who did not return for the second visit due to relocation) recruited from the New Haven community under a protocol approved by the Yale Human Investigation Committee. All participants provided written informed consent, had normal or corrected-to-normal vision, intact bilateral hand sensation by self-report, and no history of neurological or psychiatric disease. We excluded six sessions for inadequate eye-tracking calibration or persistent track loss, leaving n = 31 sessions in the analyses reported here.

### Apparatus and stimulus delivery

We collected data at the Yale Wu Tsai Institute (WTI) BrainWorks (New Haven, CT, USA). Tactile stimulation was provided using an 8-channel pneumotactile stimulus control system (Galileo Tactile Stimulator, Brainbox Ltd, Cardiff, UK),^17^ which gates compressed air to custom tactile actuators (tactile activation chambers, or TAC-cells) via computer-controlled solenoid valves at 1-ms timing resolution. The stimulator housing was located in the behavioral control room with the high-pressure line routed through a shielded wall penetration into the behavioral experiment room. Each of the eight channels was connected through pneumatic tubing to an acoustically-isolated, elastomer-sealed pulsatile pneumatic actuator (a TAC-cell),^18,19^ which we placed on the dorsum of a target digit (the index [2nd], middle [3rd], ring [4th], and little [5th] finger of each hand) midway between the proximal interphalangeal and metacarpophalangeal joints over the proximal phalanges, with channels 1 to 4 driving the left-hand cells and 5 to 8 the right-hand cells. The stimulation sites innervate the superior branch of the radial nerve (fingers 2, 3, and the lateral half of 4) and the ulnar nerve (finger 5 and the medial half of 4), providing bilateral innervation of the glabrous skin surface.^20,21^ The stimulation activates all four classes of cutaneous mechanoreceptor (fast-adapting type I and II, slowly adapting type I and II)^20,21^ with a brief air puff applied to these sites.^22^

E-Prime 3.0 (Psychology Software Tools, Pittsburgh, PA, USA) controlled stimulus delivery and timing, including the full experimental task with calibration, by issuing serial commands to the Galileo over USB 2.0 to open each channel’s solenoid for a specified pulse width. Per-channel BNC TTL outputs from the Galileo were recorded on a parallel-port acquisition line and used to verify stimulus timing offline. During the perceptual-threshold calibration (see below), the air-puff pulse width ranged from 0 to 60 ms in 6 ms steps. We collected participant responses to the perception and localization questions using a 4-button response box (Current Designs) connected directly to E-Prime over USB. We performed eye tracking at 1000 Hz with an EyeLink 1000 Plus (SR Research, Ottawa, ON, Canada), with the infrared illuminator and camera mounted on a desk in front of the head-stabilizing chinrest. Pupil size was recorded in ellipse mode and converted to a diameter-equivalent in millimeters before analysis.

### Threshold calibration

Participants completed a calibration phase of approximately 11 min (528 stimuli, 264 per hand) at 11 pulse widths (0, 6, 12, 18, 24, 30, 36, 42, 48, 54, 60 ms) with 6 pulses per each of 8 TAC locations, randomized with 1 to 1.5 s inter-stimulus interval, in which they pressed "1" if they felt the stimulus; no response was coded as "no." We fit the per-finger detection accuracy as a function of pulse width *x* with a Weibull psychometric function *P(x) = 1 − exp[−(x/λ)ᵏ]*, and the 50% detection threshold *x₅₀* was analytically calculated from the fitted scale parameter *λ* and shape parameter *k*. The initial operating pulse width for the first testing-phase run was set to *x₅₀*. The Weibull fit was re-estimated between runs using the running pool of threshold-trial responses, and *λ* was readjusted (holding *k* constant) to maintain 50% detection accuracy despite adaptation and fatigue.

### Per-finger stimulus adjustment

Tactile sensitivity is systematically distributed between fingers, based on anatomic differences in nerve distribution (fingers 2 and 3 are innervated only by the superior branch of the radial nerve, finger 5 only by the ulnar nerve, and finger 4 by both) and individual differences in mechanoreceptor density and subcutaneous fat.^20,21^ Prior to testing, we applied two complementary adjustments to equalize the per-finger detection probability at the 50% psychometric midpoint.

At the hardware level, the air-puff displacement was modulated by an in-house 3D-printed spacer that sat on each TAC cell and did not alter the acoustic or vibrational signature of the device, while allowing four different spacer configurations (2-, 3-, 5-, and 7-spacer stacks) to be used, with greater stack height resulting in a weaker felt stimulus for a given solenoid open duration. The experimenter chose a spacer configuration for each finger that resulted in a detection rate that fell within the dynamic range of the Weibull staircase (i.e., approximately 25% to 75% at the mid-range pulse width of the device) during pre-scan familiarization and fixed that configuration for that finger for the rest of the session.

Second, we fit the 66 pre-scan calibration stimuli (six repetitions of each of the 11 pulse widths) delivered to each finger using the Weibull psychometric function described above, and the per-finger *x₅₀* was computed and used as the operating pulse width for the first testing-phase run (with the between-run Weibull re-fit and *λ* re-adjustment (above) performed per finger). The combined hardware-and-software adjustment resulted in a per-finger threshold-detection rate of 52.0% across the full cohort within one SE of the target 50% midpoint, with no significant differences in detection rate by finger (one-way ANOVA, p = 0.31; see Figure 1).

**Figure 1.**
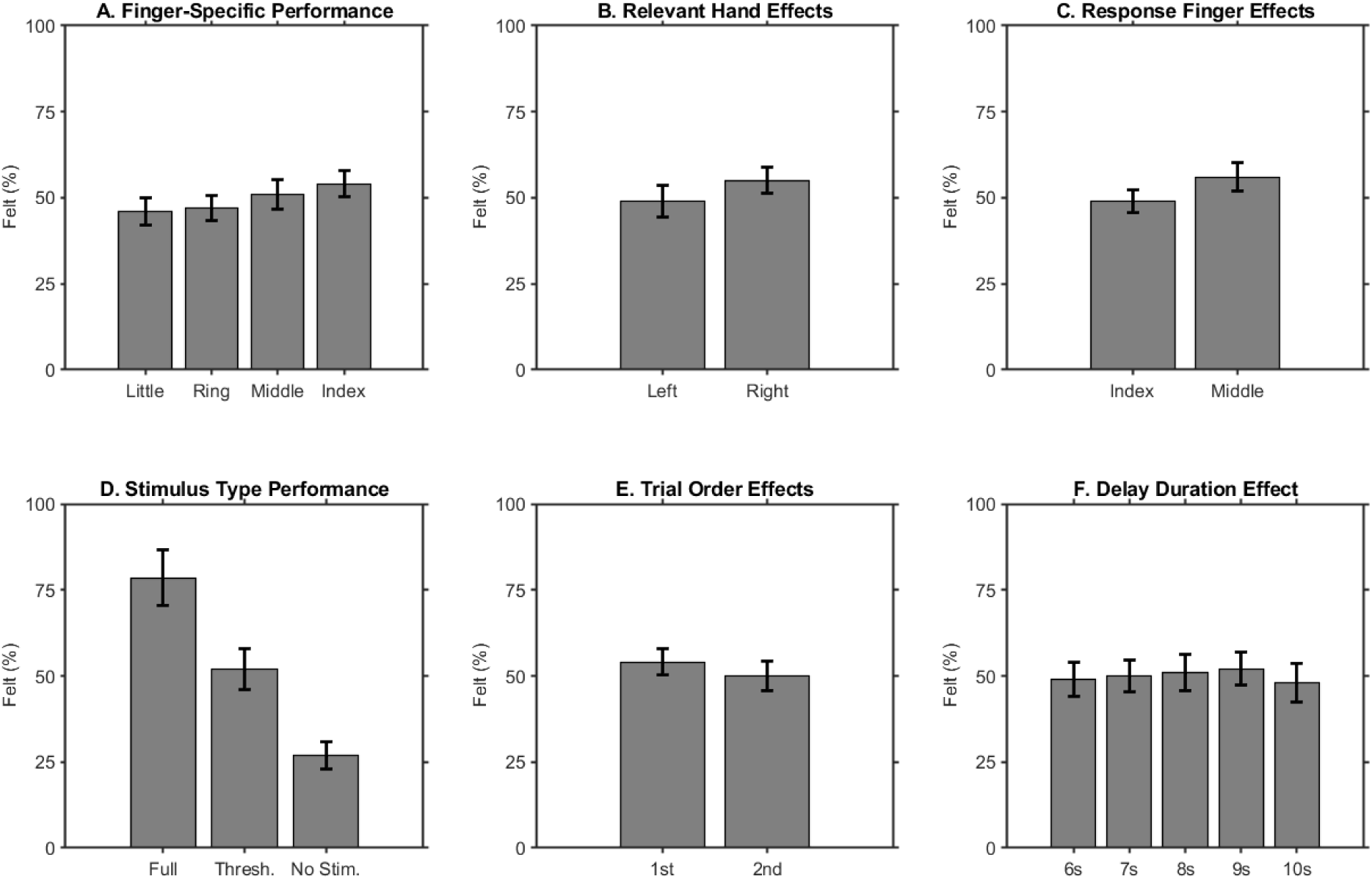
The tactile no-report paradigm is behaviorally validated. Bar graphs demonstrate the mean “felt” rate (± SEM across sessions) for (**A**) finger (little, ring, middle, index), (**B**) relevant hand (left vs. right), (**C**) response finger (index vs. middle), (**D**) stimulus intensity (Fully perceivable: 78%; Threshold: 52%; No stimulus: 27% false alarm), (**E**) trial order (1st vs. 2nd stimulus of the trial), and (**F**) delay duration (6, 7, 8, 9, 10 s inter-stimulus intervals). None of the hand, finger, trial-order, or delay-duration factors modulated the detection rate at threshold (all p > 0.3).

### Experimental paradigm

Each testing run consisted of 24 two-stimulus trials in a six-phase structure: a pre-stimulus baseline (6 to 10 s, jittered), Stimulus 1, inter-stimulus interval (6 to 10 s, jittered), Stimulus 2, post-stimulus (6 to 10 s, jittered), and a self-paced response phase. We presented three types of stimuli on both the Report and No-Report hands: Fully perceivable (12.5% of trials; well above threshold), Threshold (75%; calibrated to *x₅₀*), or Blank (12.5%). We acquired six runs (144 trials, 288 stimuli) per session, with Report and No-Report hand assignment fixed within a session. Participants were informed that one hand was task-relevant and the other was a "task-irrelevant" or "control hand" that they could largely ignore; a central fixation cross was maintained throughout. The response phase presented two questions in sequence: (1) "Did you feel a stimulus?" (yes or no) and (2) "Where was the stimulus located?" (answer choices corresponding to the 4 fingers) on the Report hand only. No questions were asked about stimuli applied to the No-Report hand.

### Eye-metric extraction

We analyzed EyeLink EDF files using in-house processing code running within the MATLAB R2023b environment. Blinks were detected using the Stublinks algorithm,^23^ which quantified a data gap as a blink if it was greater than 100 ms but less than 500 ms in duration. Pupil samples were converted to millimeters, interpolated across blinks, and low-pass filtered at 20 Hz. Microsaccades were detected using the Engbert-Kliegl algorithm^24^ at a velocity threshold of 6 standard deviations and minimum duration of 6 ms (events with amplitude < 1° or occurring within 50 ms of a blink were excluded). All four signals were epoched from −6 to +6 s around each stimulus onset and baseline-corrected using the −1000 to 0 ms pre-stimulus window.

### Single-trial classification

For each threshold stimulus, four time-locked feature streams were extracted following Kronemer et al., 2022: artifact-interpolated pupil diameter, binary blink occurrence, binary microsaccade occurrence, and artifact-interpolated gaze position (x and y), each sampled over the 12 s stimulus-centered epoch corresponding to DPSW. We then trained a stacked support vector machine (SVM) ensemble^3^ on the Report-trial threshold data for which the ground-truth perception label was available due to participant responses. The ensemble consisted of three base learners: an ensemble boosted decision tree, a linear-kernel SVM, and a Gaussian-kernel SVM ^3^(see Kronemer et al., 2022 Supplementary Figure 3). We trained each base learner independently for each feature, then concatenated their twelve output scores. A linear SVM meta-classifier combined the stacked scores to generate a final decision score in the interval [−1, +1]. Only trials with |score| > 0.75 were labeled with a confident label: trials with a score above +0.75 were labeled Predicted-Perceived, trials with a score below −0.75 were labeled Predicted-Not-Perceived, and trials in the interior band (±0.75) were considered unlabeled and excluded. Classifier performance was assessed using 10-fold cross-validation^3^ and permutation testing. We then applied the trained model without retraining to the No-Report threshold trials from the same sessions.

### Statistics

We calculated mean time courses for pupil size, blinks, and microsaccades separately for perceived and not-perceived threshold trials. We then calculated the Perceived – (minus) Not Perceived difference for each session and tested each time point against zero with a one-sample t-test (group-level α = 0.05) across the 0-to-+5-s post-stimulus window. We defined clusters as runs of consecutive samples exceeding the cluster-forming threshold (p < 0.05) and tested them against a null distribution of maximum cluster masses derived from 5000 sign-flip permutations (cluster-level α = 0.05).

## Results

### Behavioral threshold calibration generalizes across hand, finger, and trial context

For the Report hand, threshold stimuli were detected 52.0% of the time, confirming that the Weibull-fit calibration achieved the intended psychometric midpoint (Figure 1). Fully perceivable stimuli were detected 78.3% of the time, and blank stimuli were correctly rejected 73.4% of the time. The threshold detection rate did not vary by hand (left vs. right, p = 0.42), finger (one-way ANOVA across the index, middle, ring, and little fingers, p = 0.31), run position (first vs. last run of the session, p = 0.58), or inter-stimulus interval length (6, 7, 8, 9, 10 s, p = 0.49). The 50% titration generalized across the required dimensions, producing a clear within-subject perceived-versus-not-perceived contrast for physically identical stimuli and indicating the paradigm’s internal validity.

### Pupil dilates for perceived tactile stimuli (DPSW Pulse marker)

The pupil diameter time-locked to the onset of stimulation, plotted separately for perceived and non-perceived outcomes, differed significantly in the first second after stimulation (Figure 2A, top): perceived threshold trials showed a mean dilation of +0.8 mm, peaking at approximately 1.5 s post-stimulus (cluster-corrected p < 0.001). Fully perceivable stimuli showed a larger but qualitatively identical dilation profile. Both blink rejection and baseline correction controls (which limit analysis to trials in which no blink occurs within ±500 ms of the stimulus) did not alter the effect, which is consistent with H1. Overall, this supports the idea that pupil dilation induced by the task can be used as a noninvasive marker of the LC-NE Pulse component of the DPSW.

**Figure 2.**
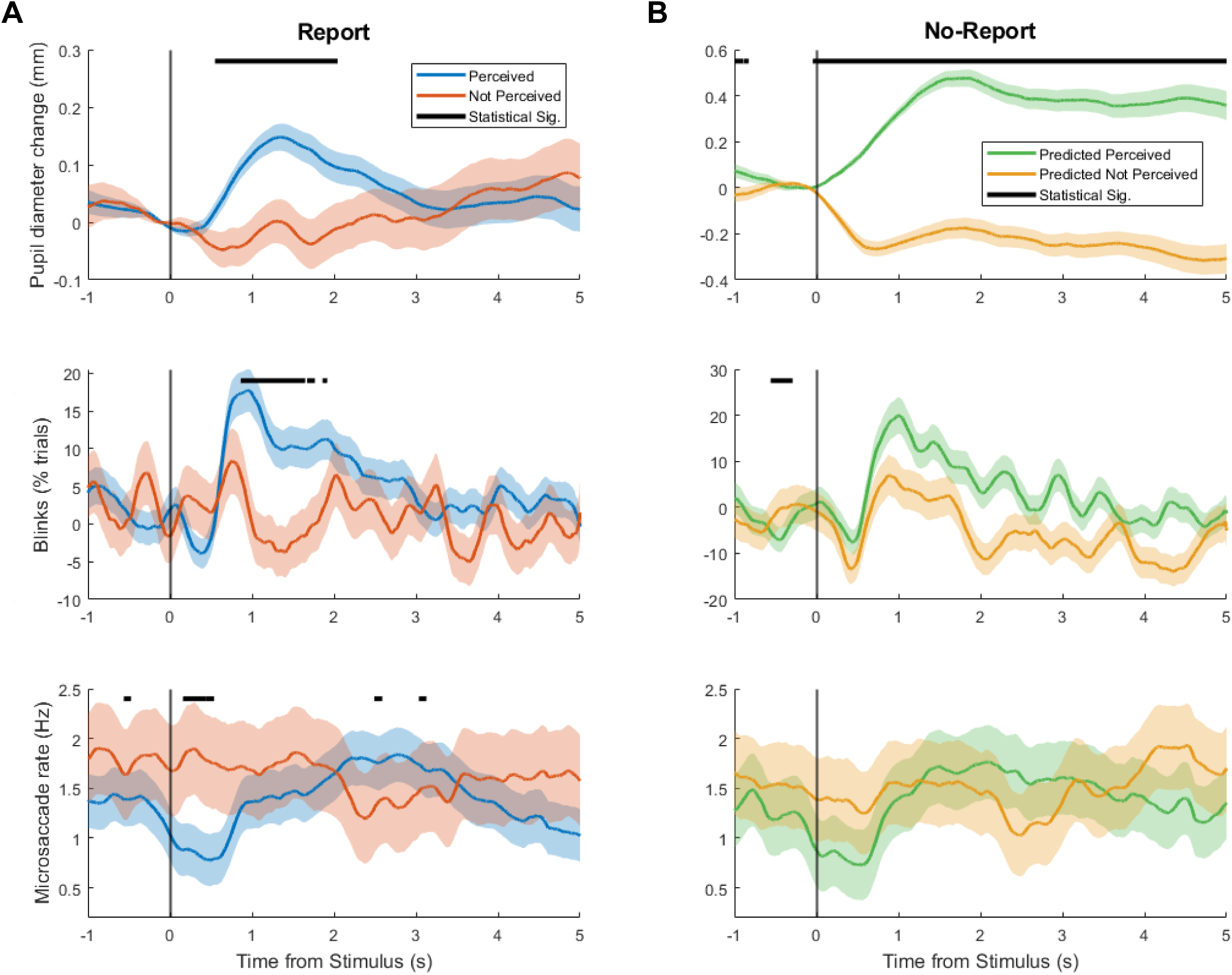
Eye-tracking signatures of conscious tactile perception in Report and No-Report trials. Eye-tracking time courses (±SEM across sessions), aligned to stimulus onset, are shown for each eye metric and colored by perceptual outcome (Perceived, blue; Not Perceived, red). Black bars mark significant between-outcome differences identified by cluster-based permutation (p < 0.05). Vertical black line marks stimulus onset. (**A**) In perceived trials, pupil dilation (baseline-corrected, in mm; top) is sustained and peaks ∼1.4 s after stimulus onset (cluster-corrected p < 0.001). Blink rate (% of trials; middle) increases at 1–2 s (cluster-corrected p < 0.001). Microsaccade rate (Hz; bottom) decreases at ∼0.5–1.5 s (cluster-corrected p = 0.08, trend). (**B**) No-Report trials were classified as Predicted Perceived (green) versus Predicted Not Perceived (yellow) using the stacked SVM ensemble without retraining (see Methods). The use of Report (A) data to train the SVM and then classify No-Report data is the central empirical result.

### Blink rate increases 1 to 2 s after perceived tactile stimuli (DPSW Switch marker)

The blink rate rose above baseline levels approximately 1 s after perceived stimuli and remained elevated for up to 2 s (Figure 2A, middle; cluster-corrected p < 0.001, 1.0 to 2.1 s). The latency and duration of the blink-rate difference are congruent with the predicted Switch window of DPSW: a neural latency of 200 to 500 ms, plus an approximate blink-execution delay of 600 to 1,000 ms. These data confirm H2.

### Microsaccade rate is suppressed around perceived tactile stimuli (DPSW Detect marker, trend)

In the perceived trials, the microsaccade rate decreased below baseline from 0.5 to 1.5 s after stimulation, whereas no such suppression occurred in the not-perceived trials (Figure 2A, bottom). The perceived minus not-perceived difference in microsaccade rate reached a corrected p-value of 0.08 at the group level. Although this result falls below the conventional threshold for statistical significance in this pilot study, it trends in the predicted direction and over the predicted timescale. These findings support H3: suppression of microsaccades by the superior colliculus is a correlate of the detection phase. Confirmation will require a larger cohort, which is currently being recruited.

### Combined eye metrics decode conscious tactile perception at AUC = 0.92

Each feature decoded trial-by-trial perception on Report trials with moderate accuracy (pupil AUC = 0.58; blink = 0.52; microsaccade = 0.43; chance = 0.50). The stacked SVM ensemble combining all four features achieved an AUC of 0.92, with 72.3% cross-validated classification accuracy at the ±0.75 score threshold and 90.1% sensitivity at 80% specificity (label-permutation p < 0.001). We then applied the same classifier, without retraining, to No-Report threshold trials from the same session. It classified 44.2% of these trials as Predicted-Perceived or Predicted-Not-Perceived; the remaining 55.8% fell within the unlabeled band and were excluded. This transfer from Report trials, where behavioral responses provide ground truth, to No-Report trials shows that the eye-based signature of conscious perception identified on Report trials is also present on No-Report trials from the same session, without an overt response.

## Discussion

Here we report that three peripheral oculomotor and autonomic signals: pupil dilation, blink rate, and microsaccade rate exhibited a strong time-locked graded pattern (pupil dilation [p < 0.001] and blink rate [p < 0.001] differed between perceived vs physically identical not-perceived threshold stimuli; microsaccade rate differed in the predicted direction with trend-level cluster-corrected p-value of 0.08) for conscious tactile perception, in a dataset consisting of out-of-scanner datasets collected from nineteen participants (38 total). We trained an ensemble model based on four features via combined stacked SVM, which resulted in an AUC value of 0.92 when predicting held-out Report trials. The same model generalized without retraining to No-Report trials during these sessions, correctly classifying approximately 44% of No-Report trials that had been previously identified as such by explicit report with the exact same model and no additional training. These results provide non-invasive physiological evidence, based on four features trained via a combined stacked SVM, of two critical components (Pulse and Switch) from the Detect-Pulse-Switch-Wave model of consciousness.^11^

The predicted mapping of pupil dilation, blinking, and microsaccades to the Pulse, Switch, and Detect components of DPSW was supported. Each signal changed in the predicted direction and on the predicted timescale in response to physically identical stimuli that differed only in conscious perception. Peak pupil dilation occurred 1–2 s after stimulus onset, consistent with the latency of task-evoked LC-NE arousal reported in human and non-human primates.^12,13^ Additionally, blink rate increased ∼1 s after stimulus onset, consistent with the predicted basal-ganglia Switch window, which includes 200 to 500 ms of neural latency and approximately 600 to 1,000 ms for blink execution. Microsaccade suppression after stimulus onset occurred as predicted, though it did not reach statistical significance in this initial pilot. However, this pattern is also consistent with better attentional control over microsaccade generation by the superior colliculus.^16^ We consider this a sample-size limitation rather than a mechanistic one; as a larger cohort is now being recruited, the predicted suppression should become significant.

The clinical relevance of this paradigm follows from its design: by using spontaneous eye movements instead of verbal reports, the paradigm could be of use for those who cannot speak, move, or follow commands^7,8^ (i.e., patients in comas or minimally conscious states, sedated or intubated ICU patients), and determining the presence or absence of conscious perception is clinically critical. Current methods of assessment rely on proxy reports, such as coma scales and behavioral evaluations,^25^ or on neural markers of arousal that do not demonstrate perception,^26^ a tactile assay that does not require cooperation or language could complement these tools and potentially be used at the bedside. The somatosensory system is especially suited to this task: it develops early, can show subcortical damage, and circumvents the imaging problems that have hampered research into somatosensory consciousness by eliminating the motor confound.

Visual no-report studies, including the seminal^3^ and its predecessors,^27,1^ have greatly advanced our understanding of the neural correlates of consciousness, but there are three reasons the tactile extension is worth pursuing. First, visual stimuli must be projected into V1 through intact oculomotor and attentional control, both of which are impaired in the clinical populations for whom no-report measurement is most needed, and tactile input through the dorsal column pathway bypasses these dependencies altogether. Second, the consciousness literature is heavily biased toward vision; we implicitly know the neural correlates of consciousness from vision, and there is weak support for generalizing to other senses.^28^ It is a nontrivial test of the model’s domain generality to show that the Pulse, Switch, and (trending) Detect signatures of DPSW appear in the tactile modality on the same timescales as in the visual modality. Third, touch is the first sense to develop and the last to fail; it is present in utero before vision has a neural substrate,^5^ and it is the only sensory modality that remains clinically active in late-stage dementia and locked-in syndrome, even when other modalities are clinically silent.^29^ Touch is thus consciousness’s earliest and most enduring signature.

These results are subject to several limitations. First, the sample size (n = 19 sessions) is sufficient for a methodological pilot but underpowered for the microsaccade effect, which was only near-significant. Second, the ±0.75 classifier confidence threshold, used for direct comparability to Kronemer et al 2022 discards around 56% of No-Report trials as unlabeled (alternative thresholds, trading precision for recall, are being considered). Third, our participants are young, healthy adults; generalization of the paradigm to the clinical populations we hope to reach will need explicit validation in each. Fourth, we have not yet replicated the findings in an independent cohort; a formal replication is underway. This preprint is meant to serve as the methodological basis for a tactile no-report fMRI study whose imaging results we will update and add when ready. The stacked SVM classifier labels approximately 44% of No-Report threshold trials with a high degree of certainty as perceived-versus-not-perceived; imaging analyses in progress (Report Confirmed Perceived − Confirmed Not-Perceived, No-Report Predicted-Perceived − Predicted-Not-Perceived, and their conjunction) will test the full set of DPSW anatomical predictions, including whether a subcortical Pulse response in the thalamus, brainstem, and basal forebrain precedes the cortical Wave in the frontoparietal task-positive network, whether the default-mode network is suppressed during the Switch, and whether these neural correlates replicate Kronemer et al.’s visual findings in the tactile modality.

Overall, these findings show that conscious tactile perception can be identified without report, pupil dilation and blink rate are effective eye-based indices of the Pulse and Switch components of DPSW in the tactile modality, and single-trial classifier labels cross over from Report to No-Report trials with sufficient consistency to support bedside clinical application. Continued work with this paradigm may offer a sensitive, non-reporting, bedside-deployable test of conscious perception in clinical populations where self-report is not feasible.

## Ethical Publication Statement

We confirm that we have followed the Journal’s ethical guidelines for publication and that this report meets those standards.

## Acknowledgments

The authors thank the Yale Electronic and Machine Shop for TAC-cell fabrication and pneumatic hardware design (Peter Brewer, Rich Rushefski); and all study participants for their time. The paradigm design builds on foundational contributions from Blumenfeld Lab alumni Sharif I. Kronemer, PhD; Wendy X. Herman, MD, PhD; and Mariana Gusso, PhD.

